# Sleep Deprivation, Sleep Fragmentation and Social Jet Lag increase temperature preference in *Drosophila*

**DOI:** 10.1101/2023.03.02.530812

**Authors:** S Tanner Roach, Melanie C Ford, Vikram Simhambhatla, Vasilios Loutrianakis, Hamza Farah, Zhaoyi Li, Erica M. Periandri, Dina Abdalla, Irene Huang, Arjan Kalra, Paul J. Shaw

## Abstract

Despite the fact that sleep deprivation substantially affects the way animals regulate their body temperature, the specific mechanisms behind this phenomenon are not well understood. In both mammals and flies, neural circuits regulating sleep and thermoregulation overlap, suggesting an interdependence that may be relevant for sleep function. To investigate this relationship further, we exposed flies to 12 h of sleep deprivation, or 48 h of sleep fragmentation and evaluated temperature preference in a thermal gradient. Flies exposed to 12 h of sleep deprivation chose warmer temperatures after sleep deprivation. Importantly, sleep fragmentation, which prevents flies from entering deeper stages of sleep, but does not activate sleep homeostatic mechanisms nor induce impairments in short-term memory also resulted in flies choosing warmer temperatures. To identify the underlying neuronal circuits, we used RNAi to knock down the receptor for *Pigment dispersing factor*, a peptide that influences circadian rhythms, temperature preference and sleep. Expressing UAS-*Pdfr*^*RNAi*^ in subsets of clock neurons prevented sleep fragmentation from increasing temperature preference. Finally, we evaluated temperature preference after flies had undergone a social jet lag protocol which is known to disrupt clock neurons. In this protocol, flies experience a 3 h light phase delay on Friday followed by a 3 h light advance on Sunday evening. Flies exposed to social jet lag exhibited an increase in temperature preference which persisted for several days. Our findings identify specific clock neurons that are modulated by sleep disruption to increase temperature preference. Moreover, our data indicate that temperature preference may be a more sensitive indicator of sleep disruption than learning and memory.

## Introduction

Although the precise function of sleep remains unknown, there is little question that sleep plays an essential role in maintaining the integrity of a large and diverse set of biological systems. In recent years, a main focus of sleep research has been on the relationship between sleep and synaptic plasticity [1; 2; 3; 4; 5; 6]. Nevertheless, it is important to note that thermoregulation has been implicated in sleep regulation and function from the earliest days of research in the field [7; 8; 9].Indeed, decades of research have firmly established that sleep and thermoregulation are inextricably intertwined on many levels [10; 11; 12]. Given this intimate interrelationship, it is exciting that recent studies have established that subsets of neural circuits regulating sleep also overlap with circuits regulating thermoregulation [13; 14; 15; 16]. However, it is unclear why the same neurons regulate sleep and thermoregulation and whether such overlap is relevant for sleep function.

The interaction between sleep and thermoregulation has clear adaptive value. That is, by coordinating the timing of sleep with daily changes in ambient temperature, animals can avoid extreme conditions and confine waking behaviors to the most optimal times of day [17]. Not surprisingly then, circadian mechanisms synergize with sleep and thermoregulatory circuits to regulate behavior. In *Drosophila*, the clock is comprised of 150 neurons that can be divided into two major groups: 1) Lateral neurons (LNd, sLNvs, lLNvs) and 2) Dorsal neurons (DN_1_, DN_2_, DN_3_) [18; 19; 20]. These neurons have been the topic of intensive investigation and are known to regulate both sleep and temperature regulation [21; 22; 23; 24]. For example, DN1s are sleep-promoting, coordinate a temperature preference rhythm and can respond to changes in ambient temperature to control the timing of sleep [15; 16; 25; 26; 27; 28]. Although clock neurons are known to regulate temperature preference across the biological day, their precise role in mediating the effects of sleep loss remains unclear.

Historically, sleep deprivation has been used as a powerful tool to evaluate sleep regulation and function [7; 29; 30]. As mentioned, sleep deprivation was used in the earliest of sleep studies where it was found that sleep loss changed temperature regulation. Importantly, sleep deprivation profoundly effects thermoregulation in rodents and humans [9; 31; 32; 33]. Indeed, feeling cold is an extremely common experience that people report during sleep deprivation [34]. Rats, like humans, also behave as if they feel cold following sleep loss. In fact, rats immediately increase operant responses for heat during sleep deprivation, indicating that changes in thermoregulation are amongst the earliest detectable changes induced by sleep loss [32]. To determine whether the effects of sleep deprivation on temperature regulation are evolutionarily conserved, we evaluated temperature preference in flies following sleep deprivation and sleep fragmentation.

## Methods

### Flies

Flies were cultured at 25°C with 50-60% relative humidity and kept on a diet of yeast, dark corn syrup and agar under a 12-hour light:12-hour dark cycle. *tim-GAL4* (BL *#7126)*; *Clk*.*856-GAL4*(BL #93198); *Clk4*.*1M-GAL4* (BL #36316); *Clk4*.*5F-GAL4* (BL #37526) and *UAS-Pdfr-RNAi* (BL #42508) were obtained from the Bloomington stock center. *per*^*01*^, *Pdfr*^*503*^, *C929-GAL4;* and *R6-GAL4* were *a* kind gift from Paul Taghert (Washington University in St. Louis)

### Thermal Preference

A 30 cm X 2.5 cm aluminum runway was positioned between a hot plate and ice to generate a gradient ranging between 30°C and 17.9°C (Figure 1A-B). Clear Plexiglas walls and ceiling were coated with Fluon (Fisher Scientific #NC0533515) to prevent flies from avoiding contact with the runway by walking on the walls. The chamber was allowed to reach equilibrium for 10 minutes and the temperature range was verified using an infrared thermometer (Fisher Scientific, 15-077-968). (Figure 1C).Thermal preference was assessed in a room at approximately 25 °C, a relative humidity of ∼50%–65% using stable light illumination. Individual flies were then introduced into the chamber for the specified time and their precise temperature was recorded using the infrared thermometer. The temperature range of the thermal gradient was validated between flies. All experiments were replicated by at least two independent investigators.

**Figure. 1.**
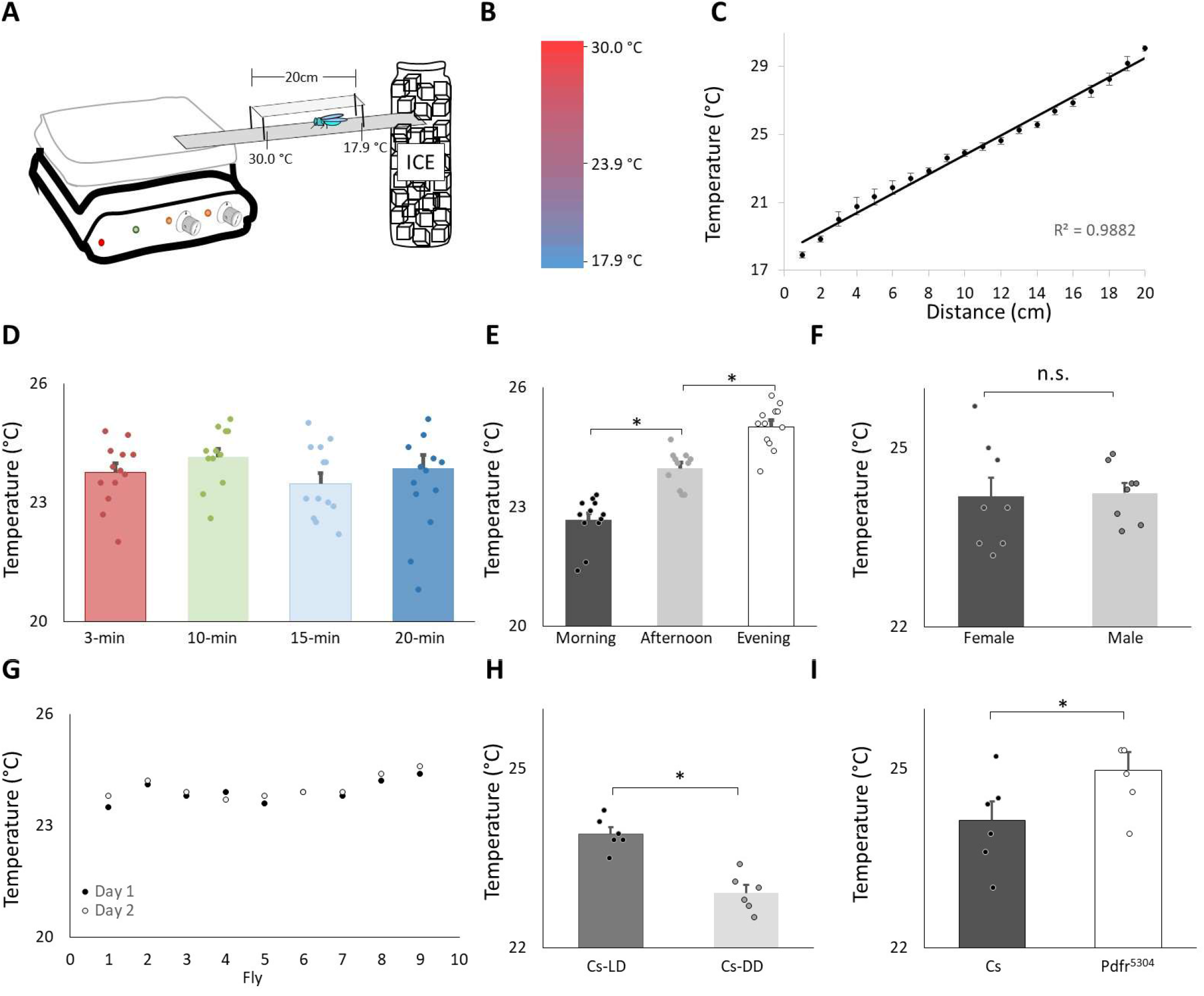
Validation of Thermal Preference Assay. **A)** Schematic for thermal preference apparatus. **B)** Heat map of the floor surface. **C)** Temperature readings in 1 cm increments measured using an infrared Thermometer (4 independent samples were taken at each location). **D)** Temperature preference in Canton-S (Cs) flies after being placed into the apparatus for different time intervals (n=11 flies/interval); One way ANOVA: F_[3,43]_ = 1.36; p=0.26. **E)** Temperature preference for Cs flies tested in the morning 8:30-10:30am, n=12; afternoon 24pm, n=11; evening 6-8pm; One way ANOVA for condition: F_[2,34]_ = 57.4; p=2.6^E-11^ *p<0.05, corrected Bonferroni test. Power analysis calculates of Cohen’s D of 8.28 between morning and afternoon. **F)** Thermal preference in male and female flies (n=8 flies/condition; p=0.86, ttest). **G)** Temperature preference in individual flies test on two consecutive days. **H)** Temperature preference in *Cs* flies tested in flies maintained on a 12:12 Light:Dark schedule (LD) and constant darkness (DD) (n=6/condition, p=9.72^E-05^, ttest). Power analysis calculates of Cohen’s D of 8.08. I) Temperature preference in *Cs* and *Pdfr*^*5304*^mutants (n=6/condition p=0.03, ttest).

### Sleep

Sleep was assessed as previously described [35]. Briefly, flies were placed into individual 65 mm tubes containing the same food as they were reared on. All activity was continuously measured through the Trikinetics Drosophila Activity Monitoring System (www.Trikinetics.com, Waltham, MA). Locomotor activity was measured in 1-minute bins and sleep was defined as periods of quiescence lasting at least 5 minutes. All sleep experiments were replicated a minimum of two times.

### Sleep Deprivation/Restriction

Sleep deprivation was performed as previously described [36; 37]. Briefly, flies were placed into individual 65 mm tubes and the sleep-nullifying apparatus (SNAP) was used to sleep deprive or sleep restrict flies Sleep deprivation was performed for 12 hours during the dark phase (lights out to lights on). For sleep deprivation, the SNAP was activated once every 20 s for the duration of the experiment. Sleep restriction was performed for 48 h. The SNAP was activated for 18 s once every 15 min for 48 hours, yielding a total of 192 stimuli lasting ∼60 minutes; this regime both reduced and fragmented sleep as previously described [38]. To determine whether the stimulus induced by the SNAP was able to alter temperature preference flies were continually exposed to the SNAP for an equal number of exposures (Bang-Control). Sleep homeostasis was calculated for each individual as a ratio of the minutes of sleep gained above baseline during the 48 h of recovery divided by the total min of sleep lost during 12 h of sleep deprivation.

### Short-term Memory

Short-term memory (STM) was assessed by Aversive Phototaxic Suppression (APS) as previously described [36; 39]. The experimenters were blinded to conditions. In the APS, flies are individually placed in a T-maze and allowed to choose between a lighted and darkened chamber over 16 trials. Flies that do not display phototaxis during the first block of 4 trials are excluded from further analysis [39; 40]. During 16 trials, flies learn to avoid the lighted chamber that is paired with an aversive stimulus (quinine/ humidity). The performance index is calculated as the percentage of times the fly chooses the dark vial during the last 4 trials of the 16-trial test. In the absence of quinine, where no learning is possible, it is common to observe flies choosing the dark vial once during the last 4 trials in Block 4 [39]. In contrast, flies never choose the dark vial 2 or more times during Block 4 in the absence of quinine [39]. Thus, short term memory is defined as two or more photonegative choices in Block 4. For short term memory experiments following a 12 h sleep deprivation, the deprivation continued until evaluation by the APS. Power analysis using G*Power calculates a Cohen’s d of 1.8 and indicates that 8 flies/group are needed to obtain statistical differences [39].

### Statistics

All comparisons were done using a Student’s T-test or, if appropriate, ANOVA and subsequent planned comparisons using modified Bonferroni test unless otherwise stated. Note that a significant omnibus-F is not a requirement for conducting planned comparisons [41]. All statistically different groups are defined as *P < 0.05.

## Results

### Validation of thermal preference

We evaluated temperature preference in adult *Canton-S* (*Cs*) flies using a slightly modified protocol from earlier studies [21; 42]. A schematic of the apparatus is shown in Figure 1A. A heat map of the runway floor is shown in Figure 1B. The temperature along the gradient, was measured using an infrared thermometer and is shown in Figure 1C. Importantly, the temperature, was stable, reproducible, and linear, with a slope of 0.6°C/cm, as previously described [42]. Previous studies have evaluated thermal preference in groups of 10-30 flies that are monitored for 20-30 min [27; 43; 44]. However, we prefer studying individual flies in which 1) the sleep history of that individual has been well characterized, and 2) the individual fly can be retrieved at the end of the assay and their behavior can be further evaluated [39; 45]. Interestingly, when we evaluated individual flies, it appeared as if they settled down and chose a preferred temperature much sooner than 20-30 min. Thus, we examined temperature preference in individual flies in 3 min, 10 min, 15 min and 20 min intervals. As seen in Figure 1D, temperature preference did not change across intervals ranging from 3 to 20 min when tested individually. It is worth noting that the temperature preference of individual flies defined using our protocol closely matches that reported by Sayeed and Benzer (1996).

Evaluating temperature preference in individual flies for 3 min differs from previous studies. Thus, we asked whether we could replicate published findings with our approach. Previous studies indicate that flies display a daily temperature preference rhythm in which they select warmer temperatures across the light period [46]. As seen in Figure 1E, when assessed after 3 min, individual flies select progressively warmer temperatures as the day progresses. Interestingly, previous studies indicate that temperature preference is similar between male and female flies run in groups. As seen in Figure 1F, no sexual dimorphisms were observed in our temperature preference assay when flies were assessed individually. To determine whether the temperature preference of an individual fly was stable, we examined temperature preference in individual flies over two successive day. As seen in Figure 1G, temperature preference is stable in an individual fly across days. A previous report indicates that flies select cooler temperature in the dark compared to siblings maintained in the light [15]. We find similar results using individual flies (Figure 1H). Finally, mutants for the receptor for the *Pigment dispersing factor receptor* (*Pdfr*^*5304*^) select warmer temperatures than controls; we find similar results when temperature preference is evaluated for 3 min in individual flies (Figure 1I). Together these data indicate that monitoring thermal preference in individual flies for 3 min accurately identifies thermal preferences and replicates published studies even when using a slightly different protocol.

### Sleep deprivation increase temperature preference

To determine the effects of sleep loss on temperature preference, *Cs* flies were exposed to 12 h of sleep deprivation. As can be seen in Figure 2A-B, sleep deprivation was effective in keeping flies awake and *Cs* flies showed a typical sleep rebound [37]. Importantly, sleep deprived flies selected warmer temperatures in the thermal gradient compared to their untreated controls (Figure 2C). We have previously shown that sleep deprivation disrupts short-term memory and that these memory deficits are completely restored after only 2 h of recovery sleep [36]. Thus, we asked how long it would take for fly’s temperature preference to return to baseline. An independent cohort of flies was sleep deprived for 12 h during their primary sleep period. The effectiveness of the sleep deprivation is shown in Figure 2D. Importantly, siblings that were allowed to recover showed sleep rebound (Figure 2E). As seen in Figure 2F, sleep deprivation resulted in flies selecting warmer temperatures replicating the data shown in Figure 2C. Similarly to what we have observed for short-term memory, 2 h of recovery sleep was sufficient to return temperature preference to baseline levels (Figure 2F). Thus, sleep deprivation increases temperature preference in flies as it does in mammals.

**Figure. 2.**
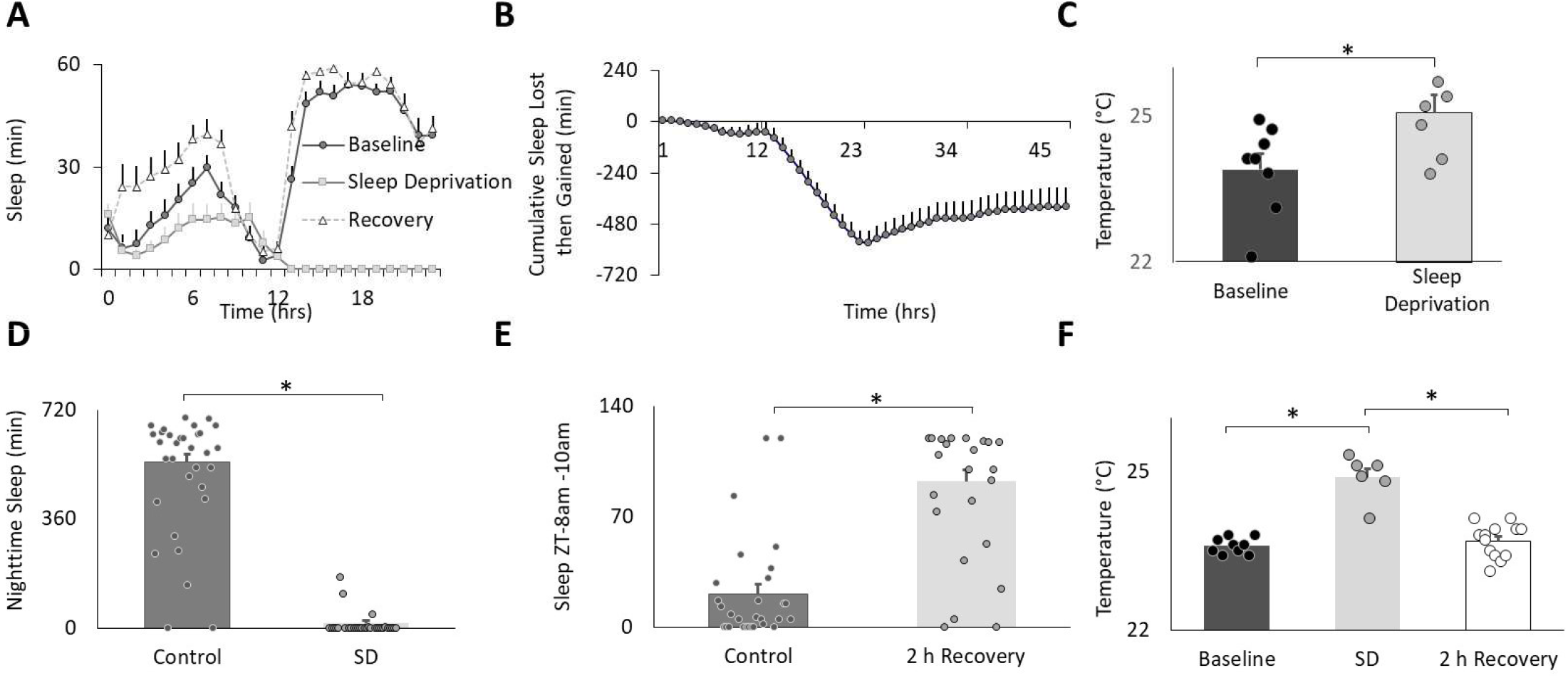
Sleep Deprivation Increases Thermal Preference. **A)** Sleep in min/hour in *Cs* flies during baseline, sleep deprivation and recovery (n=16 flies). **B)** Cumulative sleep lost then gained plot. **C)** Temperature preference is increased following 12 h of sleep deprivation (n=7) compared to baseline (n=8) (p=0.01, ttest). **D)** Nighttime sleep in untreated controls (n=30) compared to sleep deprived siblings (n=20) (p=2.40^E-22^, ttest). **E)** Sleep in untreated controls (n=29) compared to sleep deprived flies during 8-10am 2 h sleep recovery period (n=20) and (p=6.88^E-10^, ttest). **F)** Thermal preference in baseline (n=9), sleep deprivation (n=7), and sleep deprivation after 2 h of recovery sleep flies (One way ANOVA:F_[2,27]_ = 39.9; p=1.62^E-08^).

### Sleep fragmentation increase temperature preference

We have validated an ethologically relevant sleep fragmentation protocol in flies that can disrupt sleep for extended periods [38]. The protocol is based upon observations that conditions that support brain plasticity increase daytime sleep-bout duration from ∼10 min to ∼25 min [6; 36; 47; 48; 49]. These data suggest that there is a minimum amount of sleep consolidation that is required for sleep to fulfil its functions. Thus, in our sleep fragmentation protocol, flies are kept awake for 1 minute every 15 minutes to limit their ability to obtain restorative sleep. Sleep fragmentation is achieved using the sleep nullifying apparatus as previously described[38]. As seen in Figure 3A-C, sleep fragmentation modestly disrupts sleep, is effective in preventing long sleep-bouts and does not robustly activate sleep homeostasis. Importantly, sleep fragmentation did not disrupt short-term memory as assessed using Aversive Phototaxic Suppression (APS) (Figure 3D). In the APS, flies are individually placed in a T-maze and must learn to avoid a lighted chamber that is paired with an aversive stimulus (quinine/ humidity) [36]. The performance index is calculated as the percentage of times the fly chooses the dark vial during the last 4 trials of the 16 trial test and short-term memory is defined as selecting the dark vial on 2 or more occasions during Block 4 [6; 45; 50]. Despite having a normal short-term memory, flies exposed to 48 h of sleep fragmentation selected warmer temperatures in the thermal gradient compared to untreated siblings (Figure 3E). To exclude the possibility that the stimulus used to keep the animal awake altered temperature preference independently from sleep fragmentation, we exposed flies to the same number of stimuli that accrued during sleep fragmentation but in a consolidated block of 60 minutes during the light period and immediately evaluated temperature preference. As seen in Figure 3F, no changes in temperature preference were observed. Finally, we examined temperature preference during 2 h, 6 h and 24 h recovery from 48 h of sleep fragmentation. To facilitate comparisons across groups, the timing of recovery was staggered such that temperature preference was evaluated in all groups between 2:30pm and 4:30pm. This time window is ideal since flies can choose either lower or higher temperatures at this time and thereby minimizes the impact of ceiling and floor effects present during the morning and evening time points (Figure 1E)[15]. As seen in Figure 3G, temperature preference remained elevated even after 6 h of recovery. Thus, sleep fragmentation increases temperature preference in a thermal gradient and these effects persist for several hours.

**Figure. 3.**
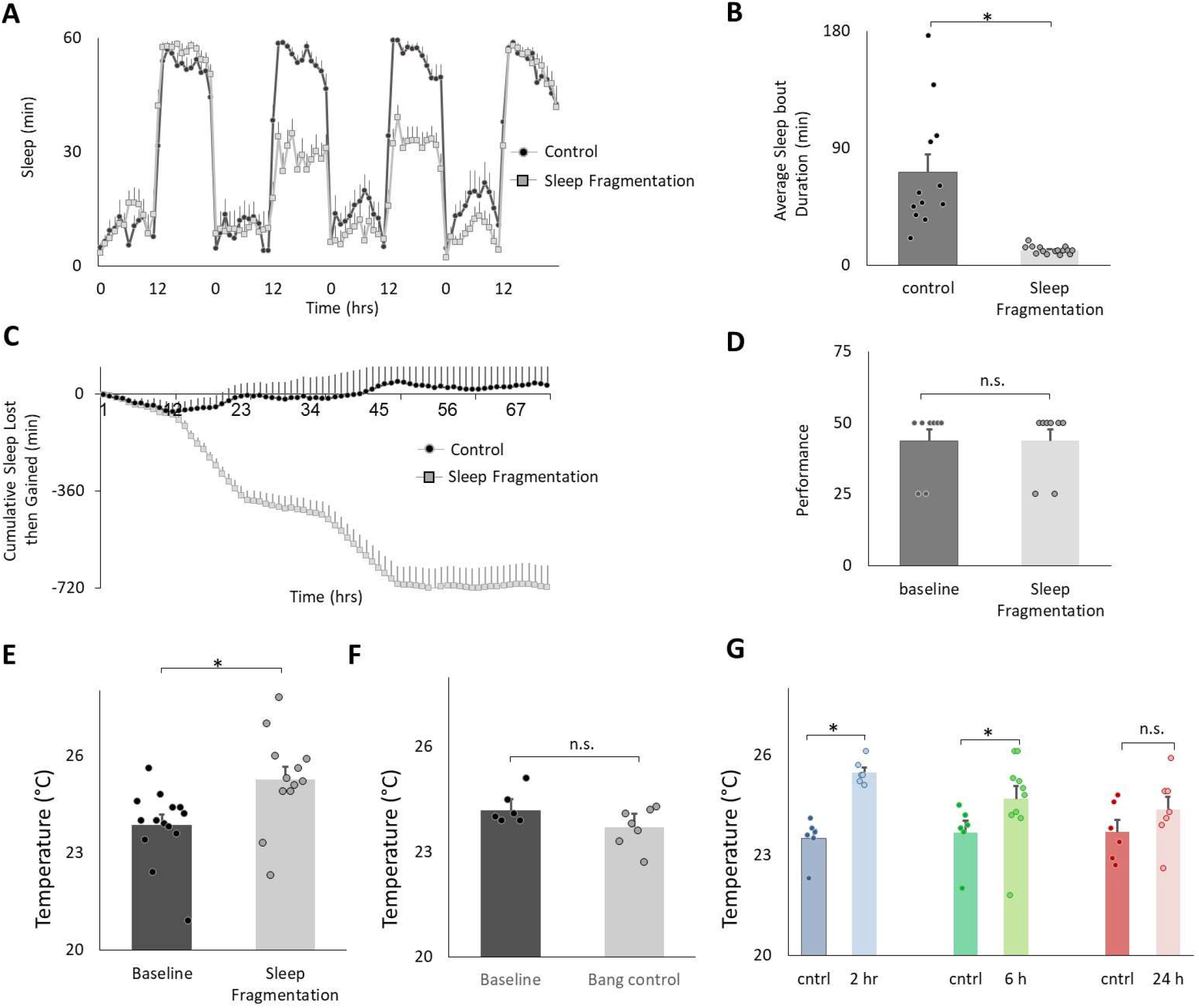
Sleep Fragmentation Increases Thermal Preference. **A)** Sleep (min/hour) in untreated controls and sleep fragmented siblings (n=12 and16 flies/condition). **B)** Average sleep bout duration during the night in untreated controls and sleep fragmented siblings (n=12 and 16 flies/condition, p=1.89^E-05^, ttest). **C)** Cumulative sleep lost then gained plot in untreated controls and their sleep fragmented siblings. **D)** Short-term memory, as assessed using Aversive Phototaxic Suppression, is not impaired by sleep fragmentation (n=8 flies/condition, p= 0.5, ttest). **E**). Sleep fragmentation increases thermal preference (n=12 and 14 flies/condition, p= 0.004, ttest). **F)** Temperature preference was not changed when flies were exposed to the same number of stimuli as sleep fragmented siblings but did not lose sleep (n=6 and 7 flies/condition, p = 0.065 ttest). **G)** Following sleep fragmentation, flies were allowed to recover for 2h, 6 h and 24 h. Thermal preference remained elevated for 2h and 6 h compared to untreated siblings. One way ANOVA F_[3,40]_ = 7.3; p=0.0005,*p<0.05, corrected Bonferroni test.

### *Pigment dispersing factor receptor* (*Pdfr*) modulates temperature preference following sleep fragmentation

Recent studies highlight the importance of the clock circuitry in regulating light dependent temperature preference [15].Specifically, light dependent temperature preference was not dependent upon the canonical clock gene *period* (*per*^*01*^) but was modulated in *Pigment dispersing factor receptor* (*Pdfr*^*5403*^) mutants. To evaluate whether these genes also play a role in the changes in temperature preference seen after sleep fragmentation, *per*^*01*^ and *Pdfr*^*5403*^ mutants were exposed to 48 h of sleep fragmentation and evaluated in the thermal gradient between 2:30 and 4:30pm as described above. As seen in Figure 4A-B, sleep fragmentation modestly disrupted sleep in *per*^*01*^ mutants without altering sleep homeostasis. Importantly, *per*^*01*^ mutants selected warmer temperature in the thermal gradient indicating that the sleep-fragmentation induced changes in temperature preference do not require the molecular clock (Figure 4C). As seen in Figure 4D-E, sleep fragmentation modestly disrupted sleep in *Pdfr*^*5403*^ mutants without altering sleep homeostasis. However, *Pdfr*^*5403*^ mutants did not select warmer temperatures in the thermal gradient (Figure 4F), suggesting that *Pdfr*^*5403*^ may play a role in mediating the effects of sleep fragmentation on temperature preference.

**Figure. 4.**
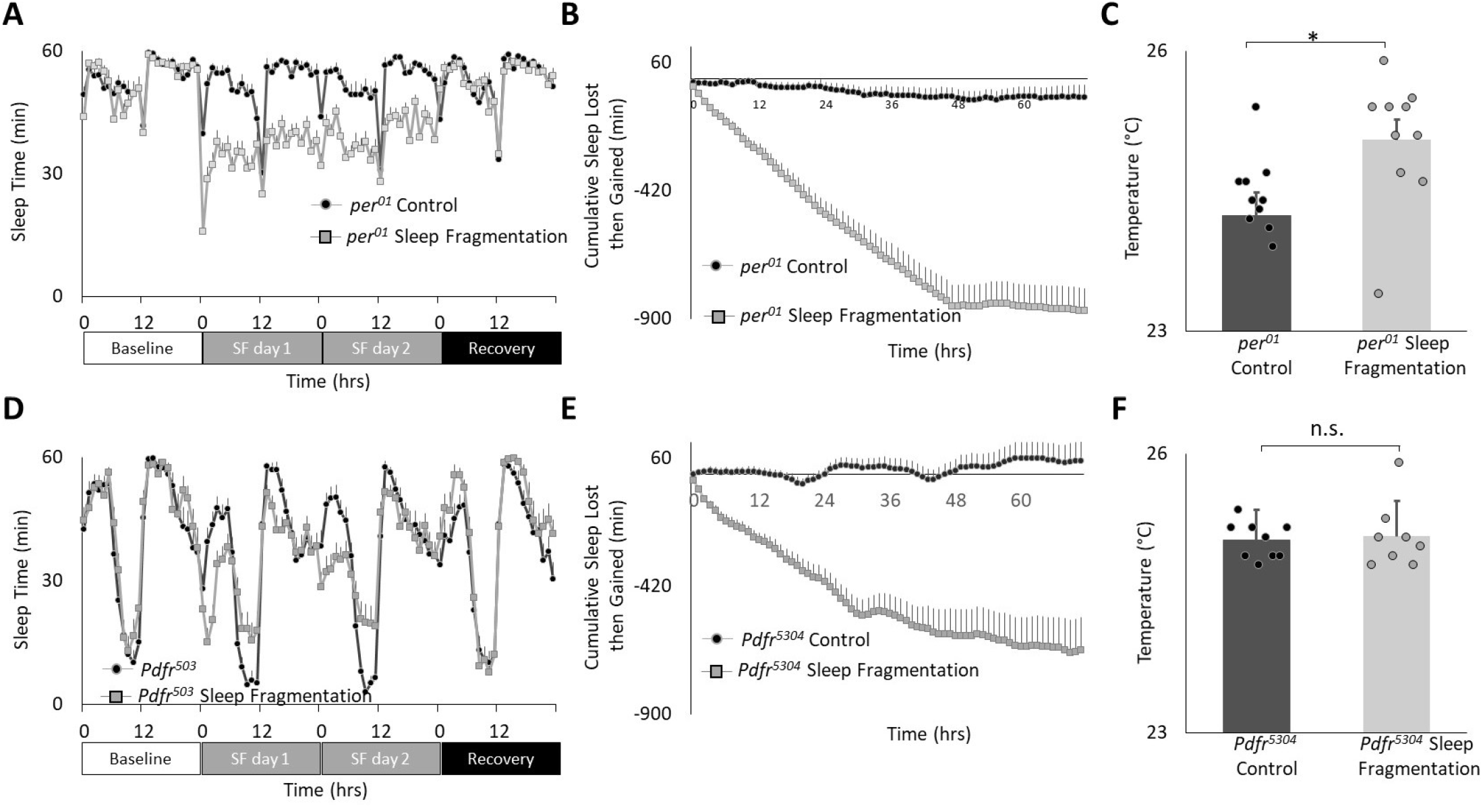
Sleep Fragmentation Alters Thermal Preference in Pdfr^5304^ mutants. **A)** Sleep (min/hour) in *per*^*01*^ mutants during baseline, sleep fragmentation and recovery (n=16 flies/condition). **B)** Cumulative sleep lost then gained plot in sleep fragmented *per*^*01*^ mutants. **C)** Temperature preference is increased in *per*^*01*^ following sleep fragmentation (n=11 and 10 flies/condition p=0. 0.01, ttest). **D)** Sleep in min/hour in *Pdfr*^*5304*^ mutants during baseline, sleep fragmentation and recovery (n=16 flies/condition). **E)** Cumulative sleep lost then gained plot in sleep fragmented *Pdfr*^*5304*^ mutants. **F)** Temperature preference is unchanged in *Pdfr*^*5304*^ following sleep fragmentation (n=7 flies/condition p=0.47, ttest)

### Clock neurons play a role in temperature preference following sleep fragmentation

The clock is comprised of 150 neurons that can be divided into two major groups. 1) Lateral neurons (LNd, sLNvs, lLNvs) and 2) Dorsal neurons (DN_1_, DN_2_, DN_3_) [18; 19; 20]. Given the role that clock neurons play in regulating temperature preference [15; 51], we evaluated their role in mediating the effects of sleep fragmentation. To begin we used RNAi to knock down *Pdfr* using the pan-clock drivers, *tim-GAL4* and *Clk856-GAL4*. As seen in Figure 5A, sleep fragmentation produced modest changes in sleep time in all genotypes. Importantly, sleep fragmentation increased temperature preference in the thermal gradient for *tim-GAL4/+, UAS-Pdfr*^*RNAi*^*/+, Clk856-GAL4/+* parental controls. However, sleep fragmentation did not alter temperature preference in the thermal gradient in either tim-*GAL4/+>UAS-Pdfr*^*RNAi*^*/+* or *Clk856-GAL4/+>UAS-Pdfr*^*RNAi*^*/+* experimental lines (Figure 5B). Together these data indicate that the clock circuitry can influence the impact of sleep fragmentation on temperature preference via the *Pdfr*.

**Figure. 5.**
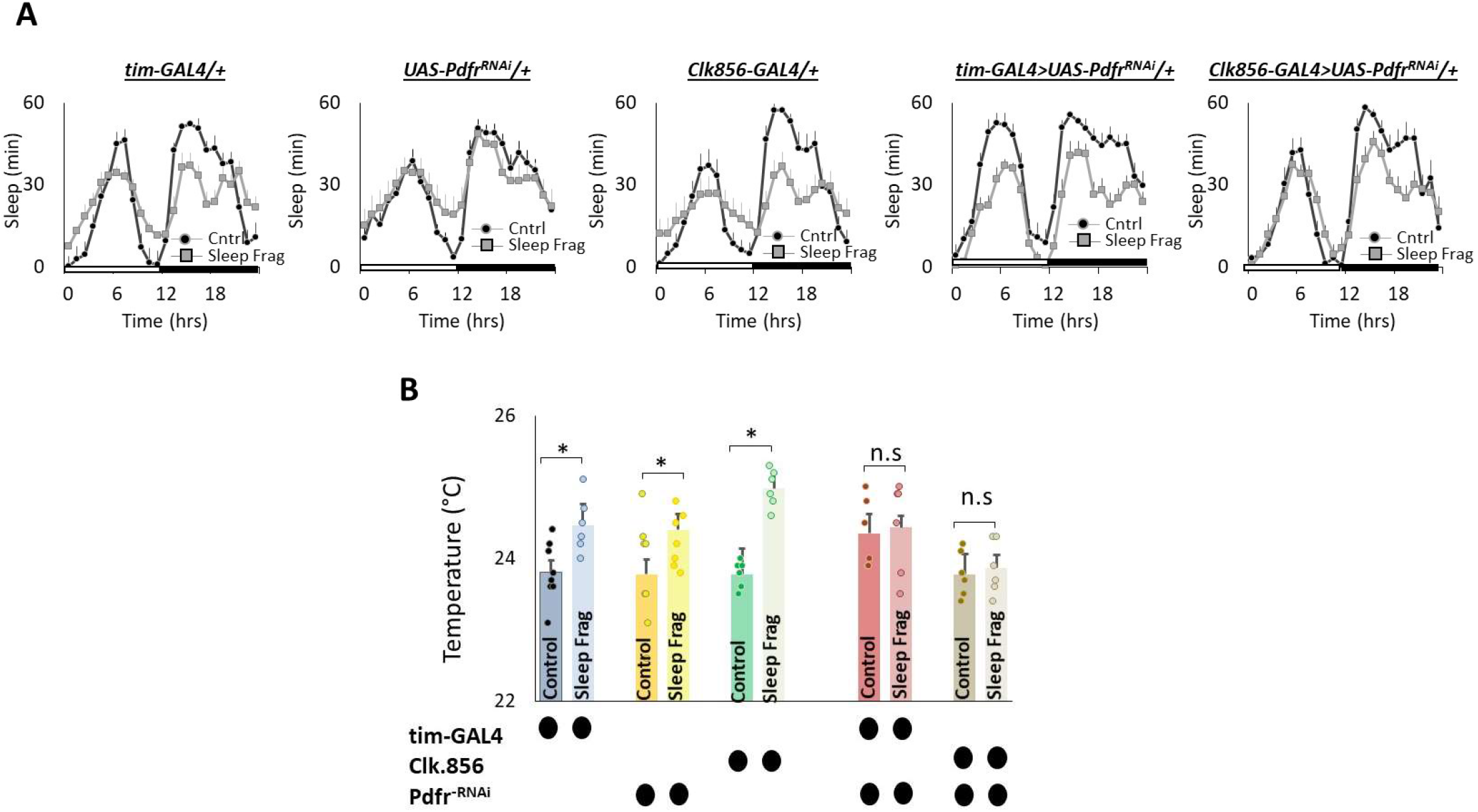
Knocking Down *Pdfr* in Clock Neurons Prevents Changes in Thermal Preference Induced by Sleep Fragmentation. **A)** Sleep in min/hour in *tim-GAL4/+, UAS-Pdfr*^*RNAi*^/+, *Clk856-GAL4/+, tim-GAL4/+*>*UAS-Pdfr*^*RNAi*^*/+and Clk856-GAL4/+>UAS-Pdfr*^*RNAi*^*/+*during baseline (Cntrl) and sleep fragmentation (n=16 flies/condition). **B)** Sleep fragmentation increases temperature preference *tim-GAL4/+, UAS-Pdfr*^*RNAi*^/+, *Clk856-GAL4/+* parental controls. Temperature preference in *tim-GAL4/+*>*UAS-Pdfr*^*RNAi*^*/+and Clk856-GAL4/+>UAS-Pdfr*^*RNAi*^*/+* is not altered by sleep fragmentation; A 5(genotype) X 2(condition)ANOVA F_[4,57]_ = 2.92; p=0.029,*p<0.05, corrected Bonferroni test.

We next evaluated the role of *Pdfr* using GAL4 lines that express in DN1 neurons. DN1 neurons play a role in sleep regulation and modulate light-dependent temperature preference [24; 25; 26; 52]. We used RNAi to knock down *Pdfr* using *Clk4*.*1M-GAL4* and *Clk4*.*5F-GAL4*. As seen in figure 6A, sleep fragmentation produced modest changes in sleep time in all genotypes. Interestingly, sleep fragmentation increased temperature preference in the thermal gradient for *Clk4*.*1M-GAL4/+*, UAS-*Pdfr*^*RNAi*^*/+* and *Clk4*.*5F-GAL4/+* parental controls. However, sleep fragmentation did not alter temperature preference in the thermal gradient in either *Clk4*.*1M-GAL4/+*>UAS-*Pdfr*^*RNAi*^*/+ or Clk4*.*5F-GAL4/+*>UAS-*Pdfr*^*RNAi*^*/+* experimental lines (Figure 6B). These data indicate that the DN1s neurons can influence the impact of sleep fragmentation on temperature preference via the *Pdfr*.

**Figure. 6.**
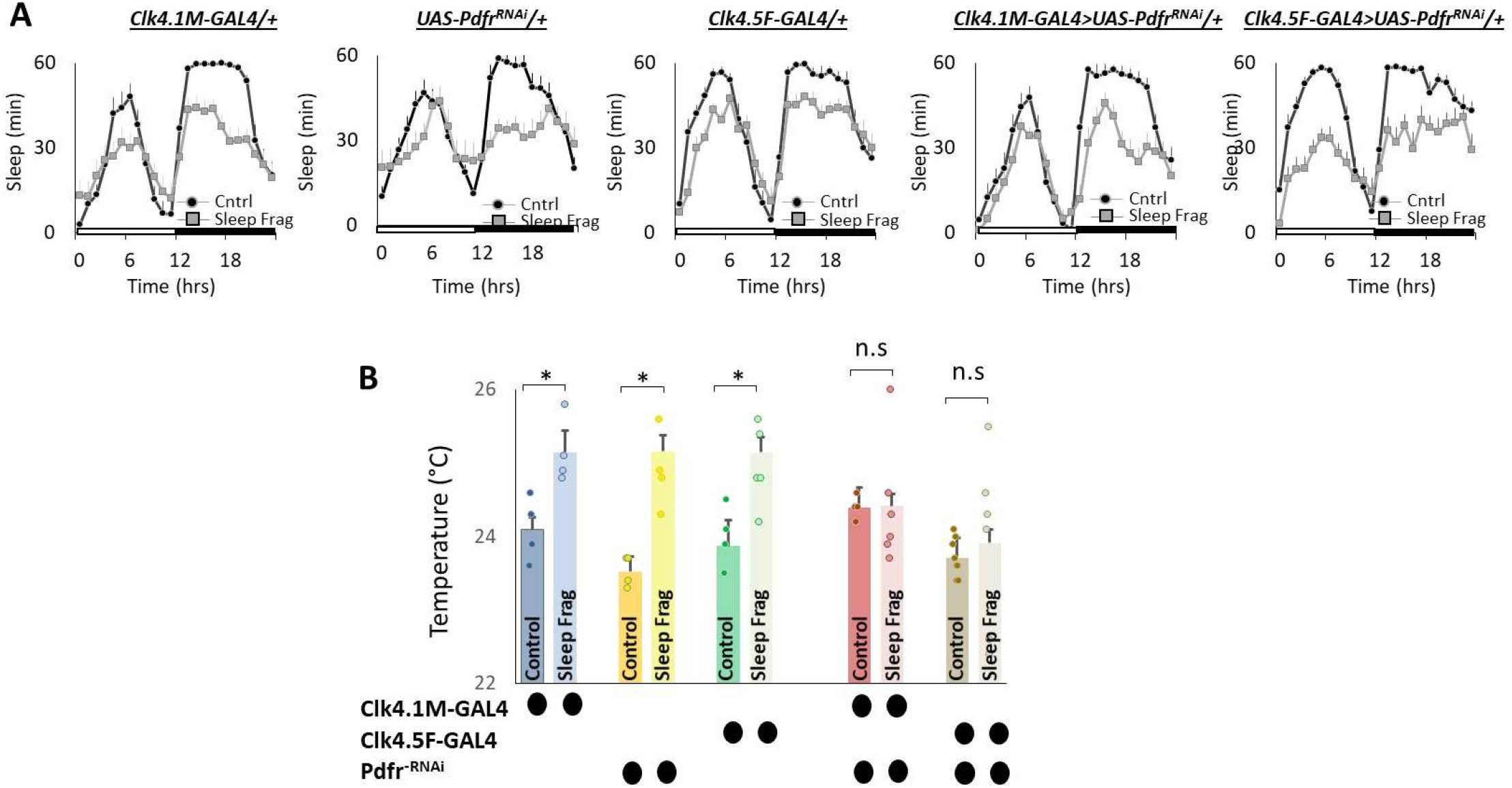
Knocking Down *Pdfr* in dorsal Clock Neurons (DN1s) neurons Prevents Changes in Thermal Preference induced by Sleep Fragmentation. **A)** Sleep in min/hour in *Clk4*.*1M-GAL4/+, UAS-Pdfr*^*RNAi*^/+, *Clk4*.*F5-GAL4/+, Clk4*.*1M/+*>*UAS-Pdfr*^*RNAi*^*/+and Clk4*.*5F-GAL4/+>UAS-Pdfr*^*RNAi*^*/+*during baseline (Cntrl) and sleep fragmentation (n=16 flies/condition). **B)** Sleep fragmentation increases temperature preference in *Clk4*.*1-GAL4/+, UAS-Pdfr*^*RNAi*^/+, *Clk4*.*F5-GAL4/+* parental controls. Temperature preference in *Clk4*.*1-GAL4/+*>*UAS-Pdfr*^*RNAi*^*/+and Clk4*.*F5-GAL4/+>UAS-Pdfr*^*RNAi*^*/+* is not altered by sleep fragmentation; A 5(genotype) X 2(condition) ANOVA F_[4,45]_ = 3.02; p=0.021,*p<0.05, corrected Bonferroni test.

Finally, we evaluated the role of the small (sLNvs) and large ventrolateral neurons (lLNvs) in modulating temperature preference after sleep fragmentation. The sLNvs and lLNvs play a role in regulating sleep and waking [53; 54; 55; 56; 57]. However, the relationship between the sLNvs and lLNvs and temperature preference is more complicated. Initial studies indicated that neither the sLNvs nor the lLNvs influence daytime temperature preference rhythms or light-dependent temperature preference [15; 46]. However, Tang and colleagues report that the sLNvs play a role in setting preferred temperature before dawn [27]. Interestingly, while pigment dispersing factor (pdf) is expressed in both the sLNvs and lLNvs; the *Pdfr* is only expressed in the sLNvs in unperturbed adult flies [58; 59]. However, the *Pdfr* is re-expressed in the lLNvs of adult flies after sleep deprivation, sleep fragmentation and starvation [38]. Since *Pdfr* is only expressed in the lLNvs of adult flies after a perturbation, its role in regulating temperature preference remains unexplored. Thus, we used *UAS-Pdfr*^*RNAi*^ to knock down the *Pdfr* in sLNvs (*R6-GAL4*) and lLNvs *(c929-GAL4*). As seen in Figure 7A, sleep fragmentation produced modest changes in sleep time in all genotypes. Interestingly, sleep fragmentation increased temperature preference in the thermal gradient for *R6-GAL4/+*, UAS-*Pdfr*^*RNAi*^*/+* and *c929-GAL4/+* parental controls (Figure 7B). However, sleep fragmentation did not alter temperature preference in the thermal gradient in either *R6-GAL4/+*>UAS-*Pdfr*^*RNAi*^*/+ or c929-GAL4/+*>UAS-*Pdfr*^*RNAi*^*/+* experimental lines. These data indicate that both the sLNvs and the lLNvs can influence the impact of sleep fragmentation on temperature preference via the *Pdfr*.

**Figure. 7.**
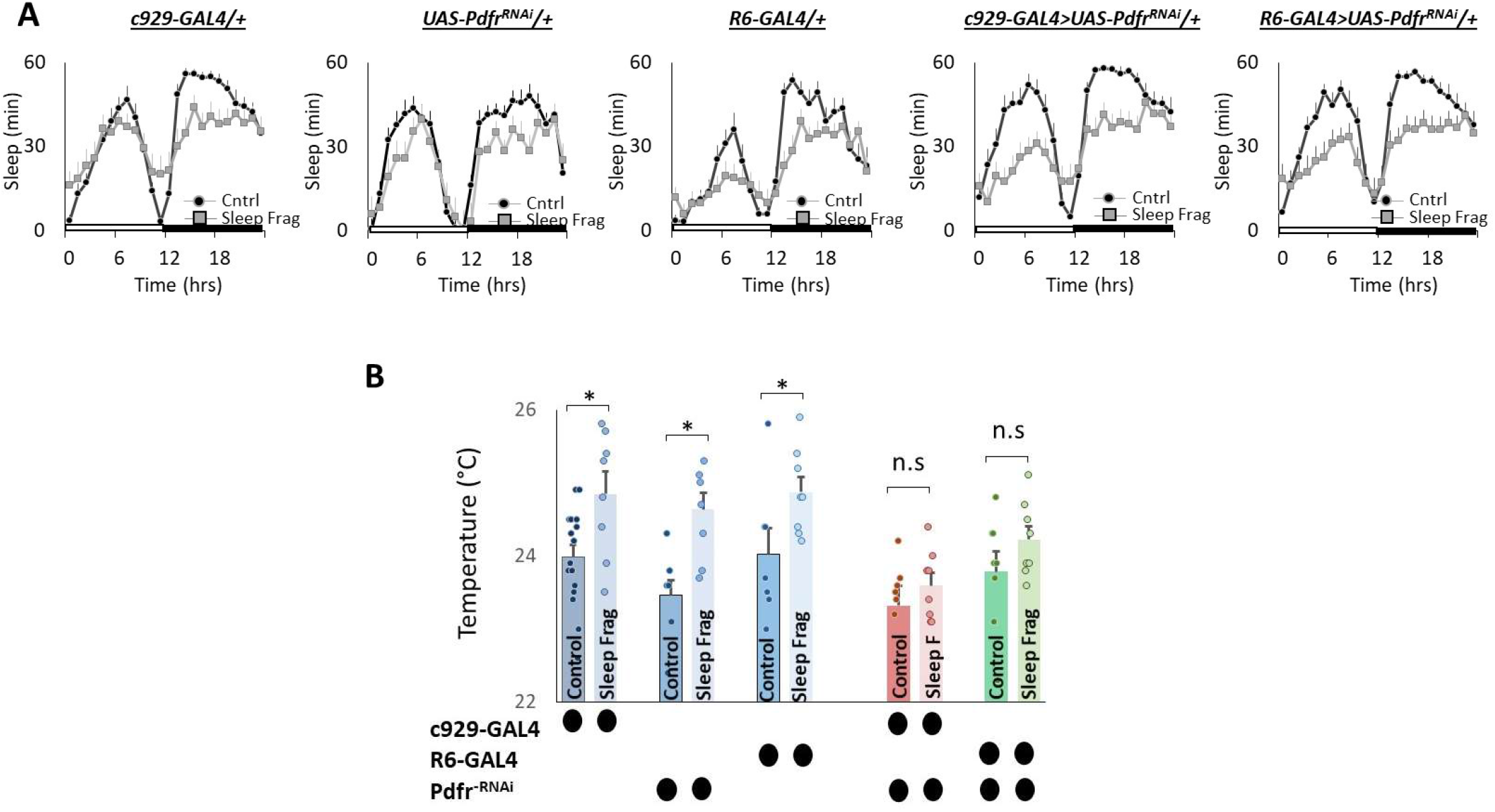
Knocking Down *Pdfr* in ventral Lateral (LNvs) neurons Prevents Changes in Thermal Preference Induced by Sleep Fragmentation. **A**) Sleep in min/hour in *c929-GAL4/+, UAS-Pdfr*^*RNAi*^/+, *R6-GAL4/+, c929/+*>*UAS-Pdfr*^*RNAi*^*/+and R6-GAL4/+>UAS-Pdfr*^*RNAi*^*/+*during baseline (Cntrl) and sleep fragmentation (n=16 flies/condition). **B**) Sleep fragmentation increases temperature preference *c929-GAL4/+, UAS-Pdfr*^*RNAi*^/+, *R6-GAL4/+* parental controls. Temperature preference in *c929-GAL4/+*>*UAS-Pdfr*^*RNAi*^*/+* and *R6-GAL4/+>UAS-Pdfr*^*RNAi*^*/+* is not altered by sleep fragmentation; A 5(genotype) X 2(condition)ANOVA F_[4,77]_ = 1.12; p=0.35,*p<0.05, corrected Bonferroni test.

### Social Jet Lag increases temperature preference

Social jet lag is a misalignment between an individual’s internal biological clock and their social schedule. Social jet lag can occur when people wake up early for work during the week and then stay up later and sleep in on the weekends to make up for lost sleep [60; 61; 62; 63]. Social Jet lag can be modeled in *Drosophila* where it has been shown to dampen rhythmicity in most circadian neurons [64]. Thus, we explored the effects of social jet lag on temperature preference in *Cs* flies. To induce social jet lag, the timing of lights-off is delayed by 3 h on Friday night and then shifted back 3 h on Sunday night to mimic a weekend schedule (Figure 8A, arrows). As can be seen in Figure 8A, sleep in flies exposed to Social Jet Lag closely resembles that seen in their age-matched untreated controls beginning on Monday after the light schedule has returned to normal to mimic a typical work week schedule. Indeed, quantification of sleep parameters, including total sleep time and sleep consolidation (average sleep bout duration during the day and average sleep bout duration at night) do not differ between flies exposed to social jet lag and their controls after being placed on a work-week schedule (Figure 8B). To further explore how social jet lag alters sleep, we evaluated metrics designed to measure sleep pressure P_Doze_ and sleep depth P_Wake_ [65]. As seen in Figure 8C, social jet lag did not alter either metric compared to untreated siblings. Social Jet lag has been reported to disrupt short-term memory [64]. We replicate that finding here (Figure 8D). In addition, to memory impairment, we show that flies exposed to social jet lag choose warmer temperatures in a thermal gradient (Figure 8D). Together these data indicate that social jet lag disrupts both learning and memory and temperature preference.

**Figure. 8.**
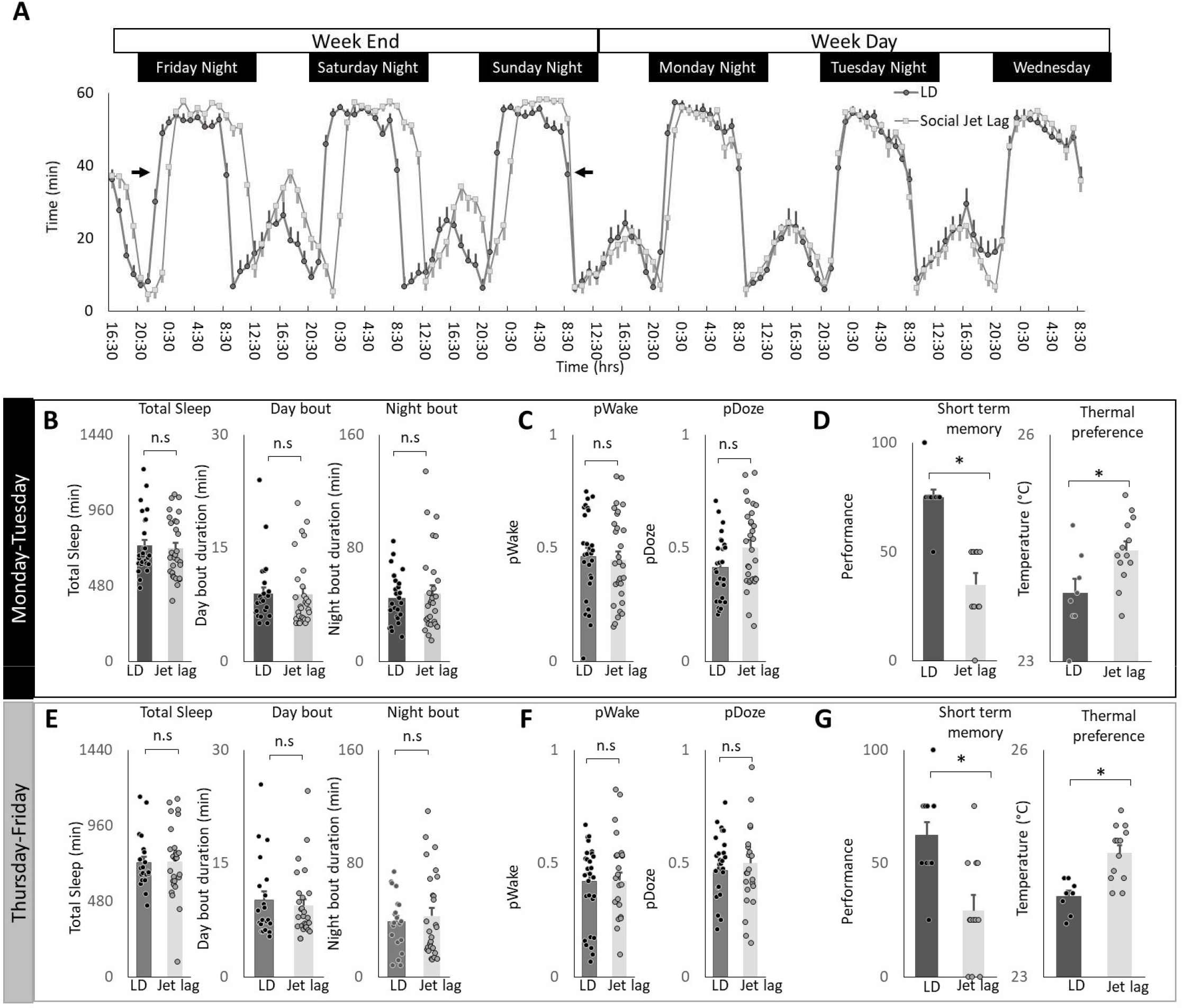
Social Jet Lag increases Thermal Preference. **A)** Sleep in min/hour in *Cs* flies maintained on a 12:12 LD schedule and siblings that have been exposed to social jet lag. For social jet lag, the timing of lights out was delayed three hours on Friday night and then advanced back three hours on Sunday night (arrows). **B)** Social Jet lag did not disrupt total sleep time, or sleep bout duration measured during the day or night compared to untreated siblings (n=28 and 30 flies/condition, p>0.05, ttest). Data are presented from Monday morning at lights-on to Tuesday morning at lights-on. **C)** Social Jet lag did not disrupt P_Wake_ or P_Doze_ (n=28,30 flies/condition, p>0.05, ttest). **D)** Social jet lag impaired short-term memory compared to untreated siblings (n= 10 flies/condition) *p=5.63^E-06^. Flies exposed to social jet lag selected warmer temperatures in the thermal gradient compared to controls (n=8 and13 flies/condition, *p=0.01, ttest). **E)** Social Jet lag did not disrupt total sleep time, or sleep bout duration measured during the day or night compared to untreated siblings (n=28 and 30 flies/condition, p>0.05, ttest). Data are presented from Thursday morning at lights-on to Friday morning at lights-on. **F)** Social Jet lag did not disrupt P_Wake_ or P_Doze_ (n=28 and 30 flies/condition, p>0.05, ttest). **G)** Social jet lag impaired short-term memory compared to untreated siblings (n= 12 flies/condition) *p= 0.001, ttest). Flies exposed to social jet lag selected warmer temperatures in the thermal gradient compared to controls (n=13 and 16 flies. condition, *p=0.0007, ttest).

To determine how long the effects of social jet lag might persist, we evaluated sleep, short-term memory and temperature preference on Friday. Recall that the light schedule had been returned to baseline on Sunday evening. As seen in Figure 8E-F measures of sleep time, sleep architecture and sleep depth are not altered by social jet lag compared to untreated controls. Despite normal sleep metrics, social jet lag treated flies shown memory impairments and increases in temperature preference several days after being placed on their typical weekday schedule. These data indicate that social jet lag results in long-lasting changes to short-term memory as well as temperature preference.

## Discussion

Our data indicate that the effects of sleep deprivation on thermoregulation are evolutionarily conserved. In addition, we show that even small disruptions in sleep induced by sleep fragmentation are able to alter thermoregulatory centers. Furthermore, we localize the effects of sleep disruption to subsets of clock neurons that play dual roles in sleep and thermoregulation. Finally, we show that while clock circuits play a role in mediating the effects of sleep disruption on temperature preference, changes in temperature preference do not require a functioning molecular clock.

The success of sleep deprivation as a tool to study sleep function depends upon the ability to monitor relevant outcome variables and their underlying neuronal circuitry. However, sleep deprivation alters a variety of physiological processes including learning and memory, metabolic, immune, digestive, cardiovascular, respiratory, and endocrine systems, to name a few [33; 66; 67]. Each of these systems are under complex regulatory control and require specialized equipment and training to evaluate properly. In contrast, temperature preference is an evolutionarily relevant outcome-variable that can be quantified using equipment found in most labs. Importantly, the temperature preference assay is exceedingly robust and the effect sizes that are seen following sleep disruption are notably high. For example, calculating the effects size for observing a difference between control and sleep deprived siblings using short-term memory reveals a Cohen’s D of 1.8, a result seen for other performance metrics in humans [39; 68]. However, when examining the impact of sleep deprivation or sleep restriction, on temperature preference the Cohen’s D is substantially higher (Cohen’s D > 3). Thus temperature preference is a unique outcome variable that can be evaluated following sleep disruption to address sleep function.

Historically, temperature preference is conducted on groups of 20-30 flies tested together as a group for 30 min. To our surprise, individual flies quickly settled down in the thermal gradient after ∼3 min. The temperatures individual flies chose are remarkably similar to the values obtained when flies were evaluated in groups [21; 42]. Importantly, we were able to replicate key findings from previously published manuscripts. Furthermore the temperature chosen by a fly was stable across days. The data presented here were collected by two independent investigators indicating that this assay produces consistent, reliable outcomes that are easily reproduced. Together, our results indicate that temperature preference in flies is a robust phenotype that can return similar results even when the protocol is changed modestly between labs.

In contrast to sleep deprivation, sleep fragmentation disrupts sleep consolidation without inducing a strong homeostatic response or deficits in short-term memory [38]. Nonetheless, sleep fragmentation is not without its consequences. Indeed, we have previously shown that sleep fragmentation can alter the expression of the *Pdfr* within the clock circuitry of adult flies. In the current study, we show that sleep fragmentation does not alter temperature preference in *Pdfr*^*5403*^ mutants or when *Pdfr* is knocked down in clock neurons. Within the clock circuitry, subsets of neurons regulate different aspects of temperature preference [22]. For example, light dependent temperature preference is mediated by DN1s, while body temperature rhythms are modulated by DN2s in concert with the sLNvs [15; 51]. Furthermore, ambient temperature feeds back on specific clock neurons to sculpt the timing of sleep and waking across the biological day [16; 28; 69]. Previous studies have shown that PDF influences the timing of behavior by differentially altering when downstream neurons display their peak activity [70; 71; 72]. Together, these observations highlight the interconnectedness of clock neurons and suggest that sleep fragmentation may disrupt temperature preference by altering the balance of activity within the clock circuit [73; 74; 75; 76].

In humans, social jet lag disrupts circadian rhythms and results in a variety of adverse health outcomes [61; 77; 78]. The social jet lag protocol we used was designed to mimic sleep/wake schedules commonly seen in humans [64]. This protocol dampens rhythmicity in most clock neurons and alters sleep regulation [64]. We show that social jet lag disrupts short-term memory and increases temperature preference. In addition, we show that the impact of social jet lag are long lasting. That is, while the light schedule is restored on Sunday evening, flies display alterations to short-term memory and temperature preference that persist at least until the following Friday. If flies were maintained on this schedule through the subsequent weekend, as is the case for many humans, they might not fully recover.

We chose to focus on PDF as a mediator of sleep loss induced changes to temperature preference given its ability to impact such a diverse set of neurons. Ongoing studies are underway to determine whether the effects of social jet lag are mediated by PDF and to identify the underlying circuitry. As noted above, PDF can modulate the timing of behavior by staggering when, during the biological day, neurons display peak activity. PDF modulates the timing of peak activity both within subsets of neurons in the clock circuit and on their downstream output targets [70; 72]. Thus social jet lag has the opportunity to disrupt short-term memory and temperature preference by disrupting a large set of diverse neurons in flies. Social jet lag has a seemingly large reach in humans as well, adversely effecting cardiovascular disease, metabolic disorders and mood. Identifying the mechanisms used by social jet lag to disrupt temperature preference in flies may provide new clues into how social jet lag adversely impacts a variety of physiological systems that are relevant for human health.

It is important to note that mechanisms regulating sleep and waking are plastic such that sleep and wake promoting neurons modulate their response properties to match sleep need with environmental demands [79; 80; 81; 82; 83; 84; 85]. For example, the wake-promoting lLNvs do not typically express the *Pdfr* in healthy adults [59]. However, the *Pdfr* is re-expressed in the lLNvs during conditions of high sleep drive, and the presence of the *Pdfr* in the lLNvs plays an important for maintaining adaptive behavior [38]. Similarly, the *Dopamine 1-like receptor 1* (*Dop1R1*) is not expressed in the sleep-promoting dorsal Fan Shaped Body neurons in healthy adults [86]. However, following starvation or time restricted feeding the *Dop1R1* is recruited to the dorsal Fan Shaped body to provide additional inhibitory tone to sleep promoting neurons [87]. Finally, a sleep circuit that is primarily active during a narrow developmental time-window can be reactivated in adults when flight is impaired [88]. These findings imply that it might be important to re-examine signaling pathways that have been previously ruled out as relevant to sleep regulation in order to better understand the effects of sleep disturbance on temperature preference. Nonetheless, understanding sleep plasticity will be important for fully understanding sleep regulation and function, and may provide crucial insight into elucidating the molecular mechanisms induced by sleep deprivation and sleep fragmentation.

The function of sleep has puzzled scientists for decades. We propose that the evolutionary origins of sleep may be better understood by determining how sleep deprivation affects neurons controlling thermoregulation in the poikilothermic fly. Both euthermic and poikilothermic animals use behavioral thermoregulation to accomplish similar goals. For example, both poikilothermic and euthermic animals can use behavior to adjust body temperature and thus conserve energy. Similarly, both poikilothermic and euthermic animals can use behavioral thermoregulation to optimize their ability to avoid predators, engage in reproductive behavior, or select the best environment for sleep [22; 89]. Indeed, ambient temperature has a profound effect on sleep in natural environments [81; 90].

Moreover, animals are known to engage in a number of behavioral adaptations that allow them to sleep in suboptimal thermal environments [89; 91; 92; 93; 94]. Thus, the interaction between sleep and behavioral thermoregulation is evolutionarily conserved. We expect that understanding the molecular mechanism linking sleep with temperature regulation in flies will provide insight into sleep regulation and function in mammals.

## Acknowledgments

We thank Matthew Thimgan, and Stephane Dissel for useful comments. We thank Paul Taghert for sharing reagents and flies. Funding: This work was supported by NIH grants 5R01NS051305-14 and 5R01NS076980-08 to PJS. The confocal facility is supported by NIH shared instrument grant S1OD21629-01A1. Competing Interests: The authors declare no competing interests.

## Notes

### Competing Interest Statement

The authors have declared no competing interest.

